# Structural Basis for Angiotensin II Type I Receptor G Protein-biased Signaling

**DOI:** 10.64898/2026.10.13.758900

**Authors:** Jinghan Liu, Meredith A. Skiba, Andrew C. Kruse

## Abstract

Biased agonism at the angiotensin II type 1 receptor (AT1R) offers a promising strategy for developing safer cardiovascular therapeutics. While β-arrestin-biased agonism has been extensively investigated, G protein-biased signaling remains poorly understood. Here, we report structures of AT1R coupled to G protein and either the endogenous agonist angiotensin II (AngII) or the G protein-biased agonist TRV055. The ligands adopt similar overall poses, with coordinated rearrangements extending from the orthosteric pocket to the intracellular surface. The structures provide a detailed view of a TM7-centered microswitch network linking ligand recognition to G protein coupling. Functional analyses of peptide analogs identify the ligand’s amino terminus as a key determinant of G protein bias, while mutagenesis of the microswitch network shifts AngII toward β-arrestin-biased signaling and converts the G protein-biased agonist TRV055 into a β-arrestin-biased ligand. These findings establish a framework for G protein-biased AT1R activation and provide mechanistic insight for the rational design of pathway-selective GPCR therapeutics.

## Introduction

G protein-coupled receptors (GPCRs) transduce extracellular signals into diverse intracellular responses and represent the largest class of therapeutic drug targets^1,2^. Upon ligand binding, GPCRs undergo conformational changes that enable coupling to multiple downstream effectors, including heterotrimeric G proteins and β-arrestins^3^. Increasing evidence suggests that distinct ligands can stabilize subsets of receptor conformations that preferentially engage specific signaling pathways, a phenomenon known as biased signaling^4–6^. This functional selectivity has generated substantial interest in drug discovery, as pathway-selective ligands may retain therapeutic efficacy while minimizing adverse effects.

The angiotensin II type 1 receptor (AT1R) is a prototypical class A GPCR that plays central roles in cardiovascular physiology and disease^7^. Around 5% of U.S. adults currently take angiotensin receptor blockers (ARBs) to manage blood pressure^8^. AT1R is activated by the endogenous peptide angiotensin II (AngII), which promotes vasoconstriction, aldosterone secretion, and cardiac remodeling primarily through Gq/11-mediated signaling^7,8^. AT1R also recruits β-arrestins, which mediate receptor desensitization and internalization while additionally initiating distinct signaling cascades^9^. The therapeutic relevance of biased signaling at AT1R has been highlighted by the development of β-arrestin-biased ligands, which were proposed to preserve cardioprotective signaling while limiting detrimental G protein activation ^9–12^. More recently, interest has also expanded to G protein-biased agonists because of their potential therapeutic applications in conditions such as septic shock and severe hypotension^13–15^. However, despite extensive pharmacological characterization of biased AT1R ligands, the structural mechanisms by which small changes in ligand sequence reshape receptor conformations and signaling outcomes remain incompletely understood.

Previous structural studies have provided initial insights into AT1R activation and signaling bias. Crystal structures stabilized by intracellular nanobodies revealed active-like receptor conformations and highlighted substantial conformational plasticity within the receptor core and TM7 region ^16^. More recently, a cryo-EM structure of an SAR1-AngII-bound AT1R–Gq complex was reported (PDB: 7F6G)^17^, providing the first visualization of G protein coupling to AT1R. However, this structure relied on a receptor construct containing two receptor-activating mutations introduced to facilitate complex stabilization and structure determination. Both mutations are known to alter receptor activation and are therefore likely to influence the observed conformations^16^. As a result, the conformational architecture and ligand interactions of wild-type AT1R in a G protein-coupled state remain incompletely defined. molecular dynamics simulations have suggested that G protein and β-arrestin biased ligands may stabilize distinct active conformations of AT1R associated with differential signaling outcomes^18^, but direct structural comparison of wild-type AT1R bound to ligands with distinct bias profiles has remained lacking.

Although modifications to the C-terminus of peptide agonists are well established as critical determinants of β-arrestin bias^12,16^, considerably less is known about how G protein-biased ligands preferentially stabilize G protein signaling. AngII and the G protein-biased ligand TRV055 differ only subtly in sequence, primarily at the ligand N-terminus, yet display markedly different signaling profiles. These observations suggest that even small changes in ligand-receptor interactions may reshape the conformational states of AT1R and bias downstream effector coupling.

Here, we report the cryo-EM structures of wild-type G protein-coupled AT1R bound to either the endogenous ligand AngII or the G protein-biased ligand TRV055. Both complexes reveal coordinated rearrangements in the receptor core and TM7 region associated with G protein coupling. Functional analyses of N-terminally modified peptide analogs further identify residues near the ligand N-terminus as important determinants of ligand efficacy and signaling bias. Together, these findings provide structural insight into the receptor rearrangements underlying G protein activation at AT1R.

## Results

### Structure determination of AT1R-Gq complexes bound to AngII and TRV055

To investigate the structural basis of ligand-dependent signaling at AT1R, we determined cryo-EM structures of the human AT1R in complex with heterotrimeric Gq, bound to both the endogenous peptide AngII and the biased agonist TRV055. To obtain a stable AT1R-Gq complex for structure determination, a miniGq chimera, miniGs/qiN^19,20^, was fused to the C-terminus of amino-terminal FLAG-tagged human AT1R, with an otherwise unmodified sequence (UniProt: P30556). An antibody fragment scFv16 was added to further stabilize the miniGs/qiN and Gβγ subunits. Complexes were formed and purified in the presence of ligands. The protein samples were then subjected to cryo-electron microscopy (cryo-EM) single-particle analysis, yielding two maps of the complex in AngII-bound and TRV055-bound at resolutions of 3.14 Å and 3.23 Å, respectively. The maps allowed unambiguous modeling of receptor, miniG protein heterotrimer, as well as the ligands (Fig. 1A-D). The interactions between AngII-AT1R and TRV055-AT1R could be clearly modeled. Structural alignments between AngII-bound and TRV055-bound receptors (Fig. 1E) showed an r.m.s.d. of 0.76 Å with well aligned intracellular regions and slightly rotations in the upper half of TM5, TM6, and TM7, particularly around the ligand-binding cavity. the overall receptor–G protein interface is highly conserved. Y374 on the Gq α5 helix hydrogen bonds with D125^3.49^ on TM3 and R140^34.57^ on ICL2 (Fig. 1E, 1F). The conserved NPxxY motif also adopts a similar active conformation in both complexes, with Y302^7.53^ in the upward position, consistent with previous molecular dynamics simulations of canonical active AT1R^18^. Hydrophobic interactions are seen between Gq α5 helix and TM3, TM5, and TM6 (Fig. 1E, 1F, and 5G).

**Figure 1.**
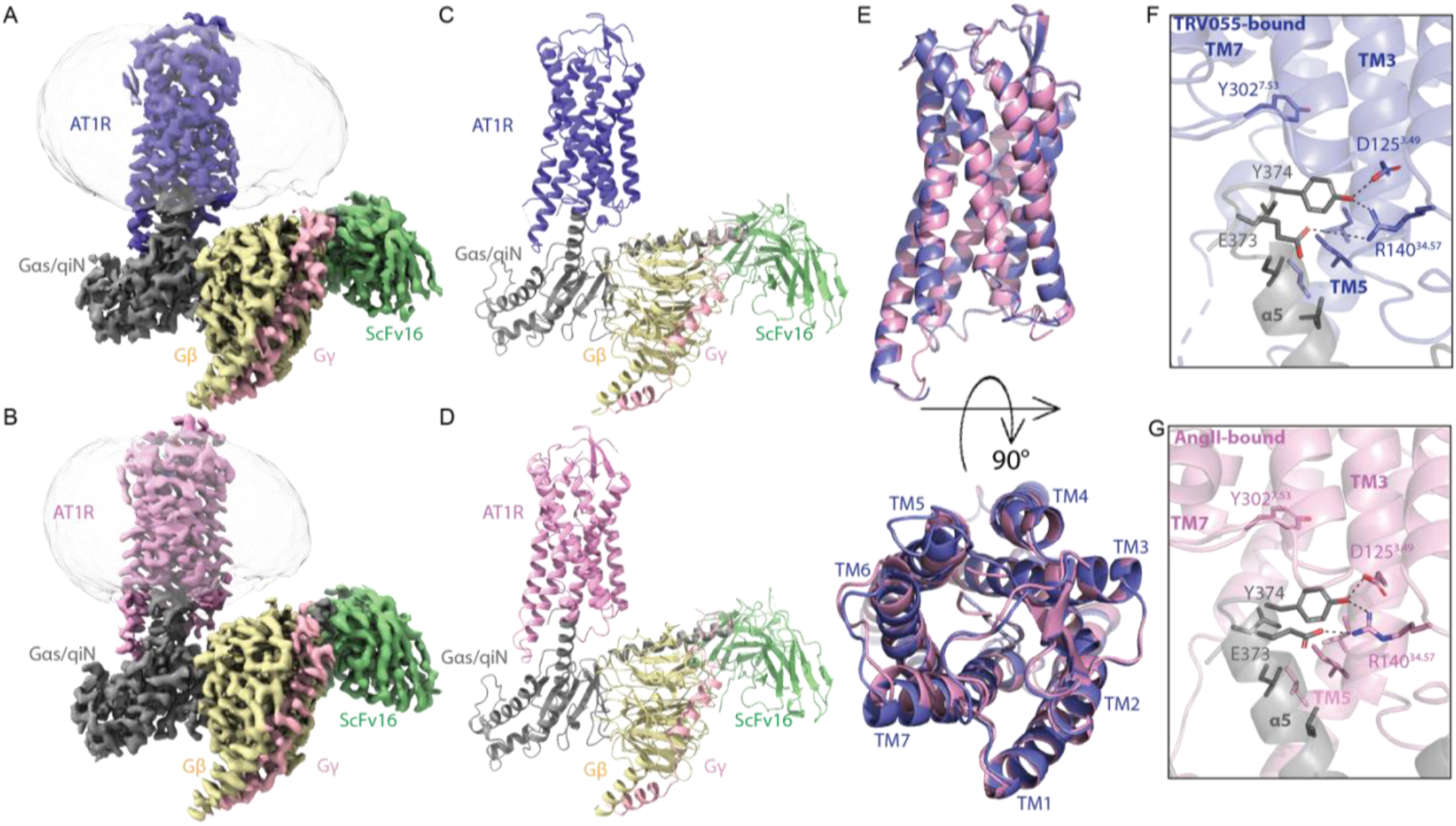
Overall structures of AT1R coupled to miniGs/q/iN. (A, C) Cryo-EM density and ribbon representation of AT1R-miniGs/q/iN-TRV055 complex, with AT1R in purple blue, Gαq in dark gray, Gβ in yellow, Gγ in light pink, and scFv16 in green. (B, D) Cryo-EM density and ribbon representation of AT1R-miniGs/q/iN-AngII complex, with AT1R in pink, Gαq in dark grey, Gβ in yellow, Gγ in light pink, and scFv16 in green. (E) Superimposition of the TRV055- and AngII-bound AT1R structure from the side and top view. (F) Interactions between TRV055-bound AT1R (purple blue) and Gq α5 helix (gray). (G) Interactions between AngII5-bound AT1R (pink) and Gq α5 helix (gray).

### Peptide analogs reveal molecular mechanisms of biased signaling at AT1R

Both AngII and TRV055 bind deeply within the orthosteric pocket of AT1R, forming an extensive interaction network spanning TM2, TM5, TM6, and TM7 (Fig. 2A, 2B). In both complexes, His6 forms a hydrogen bond with D281^7.32^, Pro7 engages in CH/π interaction with W84^2.60^, and the backbone carbonyl of Phe8 hydrogen bonds with K199^5.42^. Notably, Arg2, present only in AngII, forms a salt bridge with D263^6.58^ or D281^7.32^. This interaction pulls the extracellular end of TM7 outward while maintaining TM6 in a relatively inward position compared to the TRV055-bound complex (Fig. 2C).

**Figure 2.**
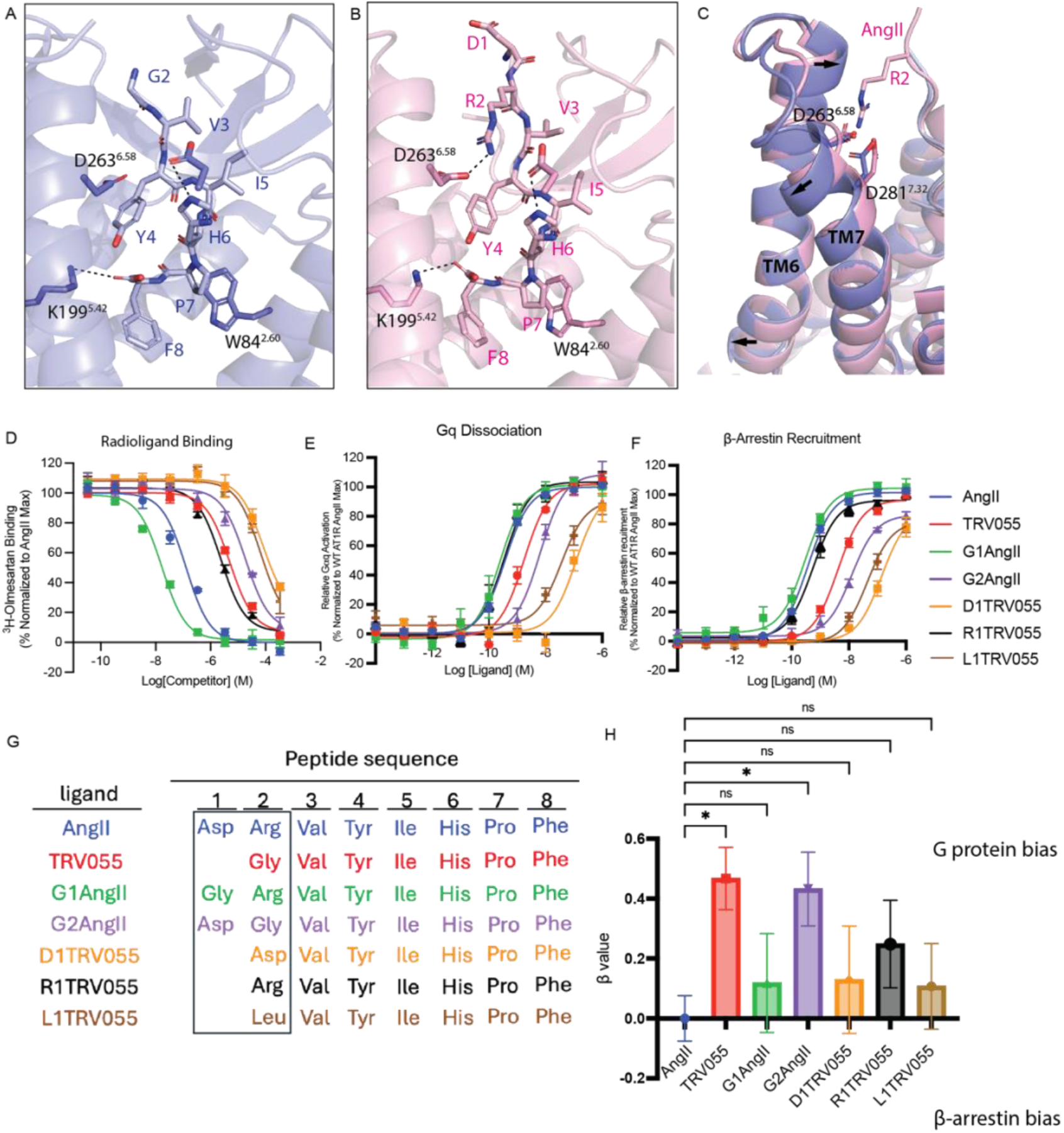
Signaling bias regulated by AngII, TRV055 and peptide analogs. (A, B), Orthosteric binding pockets of the AngII-bound (A) and TRV055-bound (B) AT1R–Gq complexes. Key ligand–receptor interactions are shown in dash. (C) Superposition of the AngII- and TRV055-bound AT1R structures viewed from the side, with TRV055-bound structure in purple blue and AngII-bound AT1R in pink. Arrows indicate conformation differences between 2 structures. (D) Competition binding between the radiolabeled antagonist [^3^H]-olmesartan and unlabeled peptides in cell membranes containing AT1R. (E, F) BRET-based Gq dissociation (E) and β-arrestin recruitment (F) assays comparing AngII, TRV055, and N-terminal peptide analogs. (G) Sequences of AngII, TRV055, and N-terminal peptide analogs used in signaling assays. (H) Bias factors calculated from Gq dissociation and β-arrestin recruitment assays. AngII was used as a reference for each ligand. Error bars represent the mean SE of at least 3 independent experiments. β values were presented as mean ± S.E.M. β > 0, Gq biased; β < 0, β-arrestin-biased. Statistics analyses were performed using one-way analysis of variance (ANOVA). P < 0.05 (*).

Given the minimal sequence differences between AngII and TRV055, we compared their signaling properties alongside a panel of N-terminally modified analogs to define how peptide sequence encodes downstream signaling outcomes (Fig. 2F). Ligand binding affinities were determined by competition radioligand-binding assays, and Gq dissociation and β-arrestin recruitment were quantified using BRET assays, from which Ki vaules and EC₅₀ values for each pathway were derived, (Fig. 2D-F). These data suggest that AngII exhibited higher binding affinity than TRV055 (logKi = -7.76 vs. -6.17) and greater potency in both Gq dissociation and β-arrestin recruitment. Notably, the difference in potency between AngII and TRV055 was substantially greater for β-arrestin recruitment (logEC₅₀ = -9.44 vs. -8.39) than for Gq dissociation (logEC₅₀ = -9.43 vs. -8.76), consistent with the G protein-biased signaling profile of TRV055. Removal of Asp1 or Arg2 in AngII produced divergent effects. G1AngII displayed increased binding affinity (logKi = -8.64) and retained high potency in both Gq (-9.61) and β-arrestin (-9.53) signaling, whereas G2AngII showed markedly reduced affinity (logKi = -5.59) and reduced potency in both pathways (-8.20 and -7.91, respectively), with a greater loss in β-arrestin signaling contributing to its increased G protein bias (0.423).

Modifications of the N-terminal Gly in TRV055 similarly produced distinct effects on affinity and potency. R1TRV055 increased binding affinity (logKi = -6.49) and potency in both Gq (-9.46) and β-arrestin (-9.27) assays. In contrast, D1TRV055 and L1TRV055 showed substantially weaker binding (logKi = -4.96 and -5.05, respectively). However, D1TRV055 retained relatively high signaling potency (-8.93 (Gq), -8.88 (β-arrestin)), whereas L1TRV055 showed pronounced losses in both pathways (-7.44 and -7.38). Despite these divergent effects on affinity and potency, substitution of the N-terminal Gly with a positively charged Arg, negatively charged Asp, or hydrophobic Leu consistently reduced G protein bias relative to TRV055 (bias factor = 0.467), with bias factors of 0.248, 0.129, and 0.107 for R1TRV055, D1TRV055, and L1TRV055, respectively, supporting the importance of the small Gly side chain in promoting preferential G protein signaling.

Together, these results indicate that the N-terminal composition of these peptides regulates signaling bias independently of overall receptor affinity and support a role for the additional space provided by Gly near D263^6.58^ and D281^7.32^ in promoting G protein-biased signaling.

### Microswitch rearrangements upon AT1R G protein activation

Despite differences at the N-terminus, TRV055 and AngII share identical C-terminal sequences and adopt similar binding poses, with Phe8 oriented toward TM7 as supported by the density map, resulting in comparable rearrangements of residues within the orthosteric pocket (Fig. 3B). Notably, the Phe8 in both AngII- and TRV055-bound complexes adopts a distinct rotamer compared to a previously reported AngII-bound AT1R crystal structure (PDB:6OS0). The AngII-bound crystal structure is stabilized by an intracellular nanobody AT110, an arrestin-mimetic nanobody that stabilizes an arrestin-coupled receptor conformation, rather than coupled to Gq heterotrimer^16^. The Phe8 was conformationally dynamic in the AT1R crystal structure, whereas, in our cryo-EM structures the TM7-directed orientation of Phe8 is well resolved without evidence of alternative conformations (Fig. S3). The differences in Phe8 conformations observed in the two structures may therefore reflect differences in the stabilized receptor state, with AT110 favoring an arrestin-coupled conformation and G protein coupling stabilizing the TM7-directed orientation.

**Figure 3.**
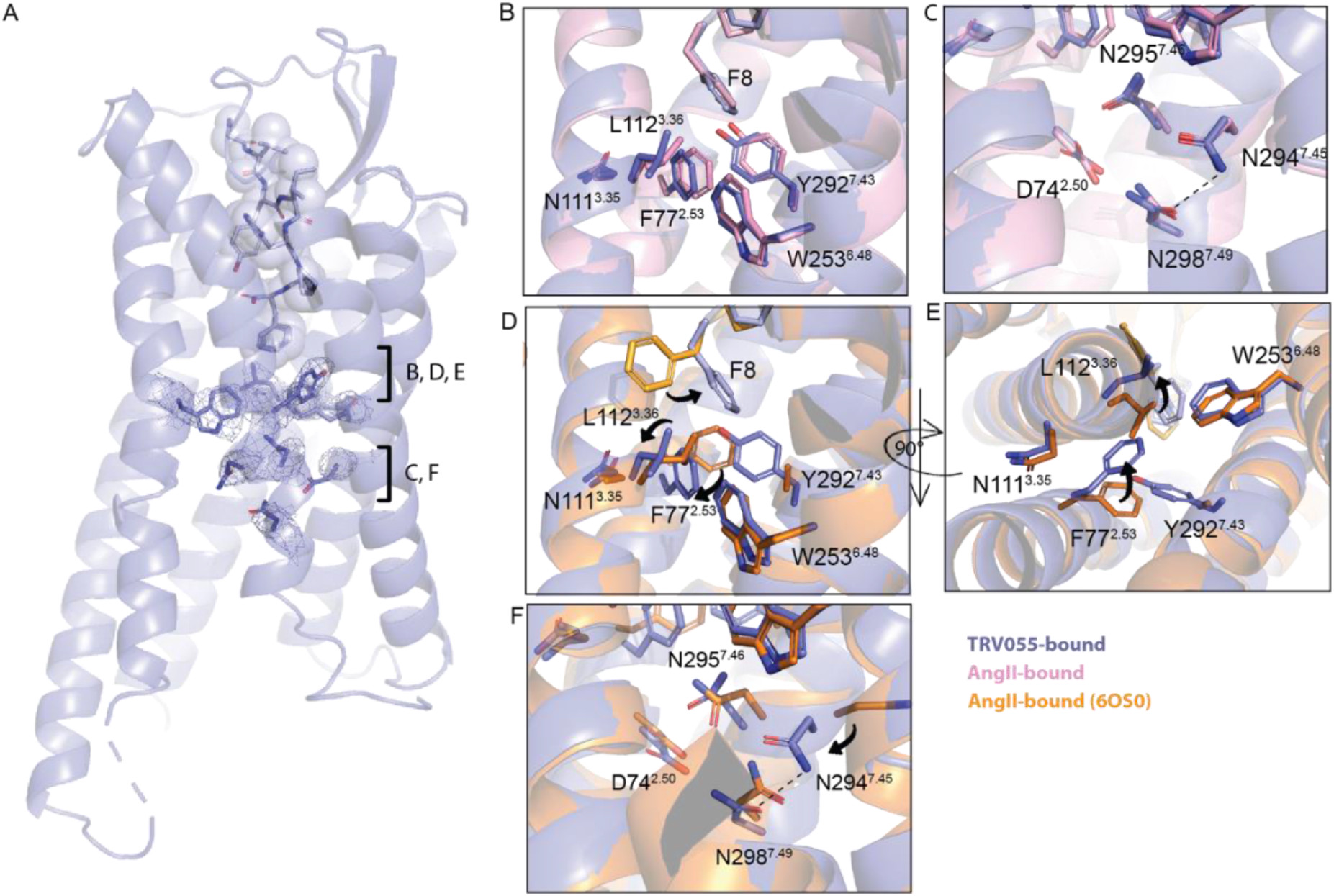
Structure comparisons between TRV055-bound AT1R and AT110-bound AngII-bound AT1R (6OS0). (A) Cryo-EM map of the ligand and key residues in the orthosteric pocket and lower TM region in the TRV055-bound AT1R–Gq complex. The boxed regions correspond to the close-up views shown in (B) and (C). (B, C) Superposition of the AngII-bound (pink) and TRV055-bound (blue) G protein-coupled AT1R structures in the orthosteric pocket (B) and lower TM region (C). (D, E) Comparison of the G protein-coupled TRV055-bound structure (blue) with the previously reported AngII-bound nanobody-stabilized AT1R structure (orange; PDB: 6OS0) at the orthosteric pocket region from side view (D) and bottom view (E). Arrows indicate conformation rearrangements in the pocket. (F) Comparison of the G protein-coupled TRV055-bound structure (blue) with the previously reported AngII-bound nanobody-stabilized AT1R structure (orange) at lower TM region. Arrow indicates conformation rearrangement.

This TM7-adjacent Phe8 pose is consistent with previous molecular dynamics simulations, which describe an “inserted” conformation in which the C-terminal Phe8 rotates toward TM7 and directly engages Y292^7.43^ ^18^. In our cryo-EM structures, this repositioning stabilizes Y292^7.43^, which is unresolved in the crystal structure (Fig. 3D), and propagates local rearrangements within the receptor core. Specifically, stabilization of Y292^7.43^ is accompanied by rotation of F77^2.53^ toward TM3 to avoid steric clash, which in turn shifts L112^3.36^ away from the binding pocket. Comparison with the antagonist-bound AT1R structure (olmesartan-bound, PDB: 4ZUD) further illustrates the extent of these rearrangements: Y292^7.43^ shifts from a TM3-directed to a TM7-adjacent conformation, accompanied by a downward shift of W253^6.48^, rotation of F77^2.53^ toward TM3, and displacement of N111^3.35^ away from the receptor core disrupting the inactive state-stabilizing interaction between N295^7.46^ and N111^3.35^ (Fig. S6 C-D). These coordinated rearrangements provide a structural link between the TM7-directed Phe8 pose and remodeling of the receptor core.

Rearrangements within the orthosteric pocket propagate to the intracellular side of the receptor, leading to coordinated changes in the lower transmembrane region. In both Gq-coupled structures, N294^7.45^ adopts a distinct conformation, rotating away from TM7 and into the receptor core, where it forms a hydrogen-bonding interaction with N298^7.49^ (Fig. 3F). Notably, a similar interaction between N294^7.45^ and N298^7.49^ is observed in the olmesartan-bound AT1R structure (PDB: 4ZUD), although N294^7.45^ adopts slightly different conformation (Fig. S6F), whereas more distinct conformations are observed in previously reported active-state structures. The presence of this interaction in both antagonist- and Gq-bound structures suggests that this interaction is not specific to a single receptor activation state, but can be accommodated within different conformational states of AT1R.

A previously reported cryo-EM structure captured AngII-bound AT1R coupled to Gq (PDB: 7F6G). To compare the active-state conformations of wild-type and engineered receptors, we superimposed our wild-type AngII-bound structure with the Sar1-AngII bound structure (PDB:7F6G) (Fig. S7). The two structures exhibit highly similar overall architectures, with the transmembrane helices and intracellular G protein interface superimposing closely. Interpretation of local differences, however, is complicated by the F77A and N295A substitutions in Sar1-AngII bound AT1R structure. The F77A substitution removes steric constraints within the orthosteric pocket and likely contributes to the distinct orientations of the C-terminal F8 and L112^3.36^ observed in the two structures. In addition, the N295A substitution disrupts the interaction between N295^7.46^ and N111^3.35^, a mutation previously shown to promote constitutive receptor activity, and likely contributes to the distinct rotamer conformation adopted by N294^7.45^ in the wild-type structure. Our wild-type structure therefore provides a view of the native active-state interaction network underlying AT1R activation in the absence of stabilizing receptor mutations.

### Distinct microswitch rearrangements upon G protein- and β-arrestin-biased ligand binding

To further investigate structural determinants associated with G protein and arresting favoring AT1R states, we compared the TRV055-bound Gq-coupled structure with a previously reported structure of AT1R bound to the β-arrestin-biased ligand TRV023. Although the overall receptor architectures are highly similar, slight helices shifts are observed in TM5, TM6, and TM7, and local conformational differences are observed within the orthosteric pocket and TM7 region (Fig. 4C).

**Figure 4.**
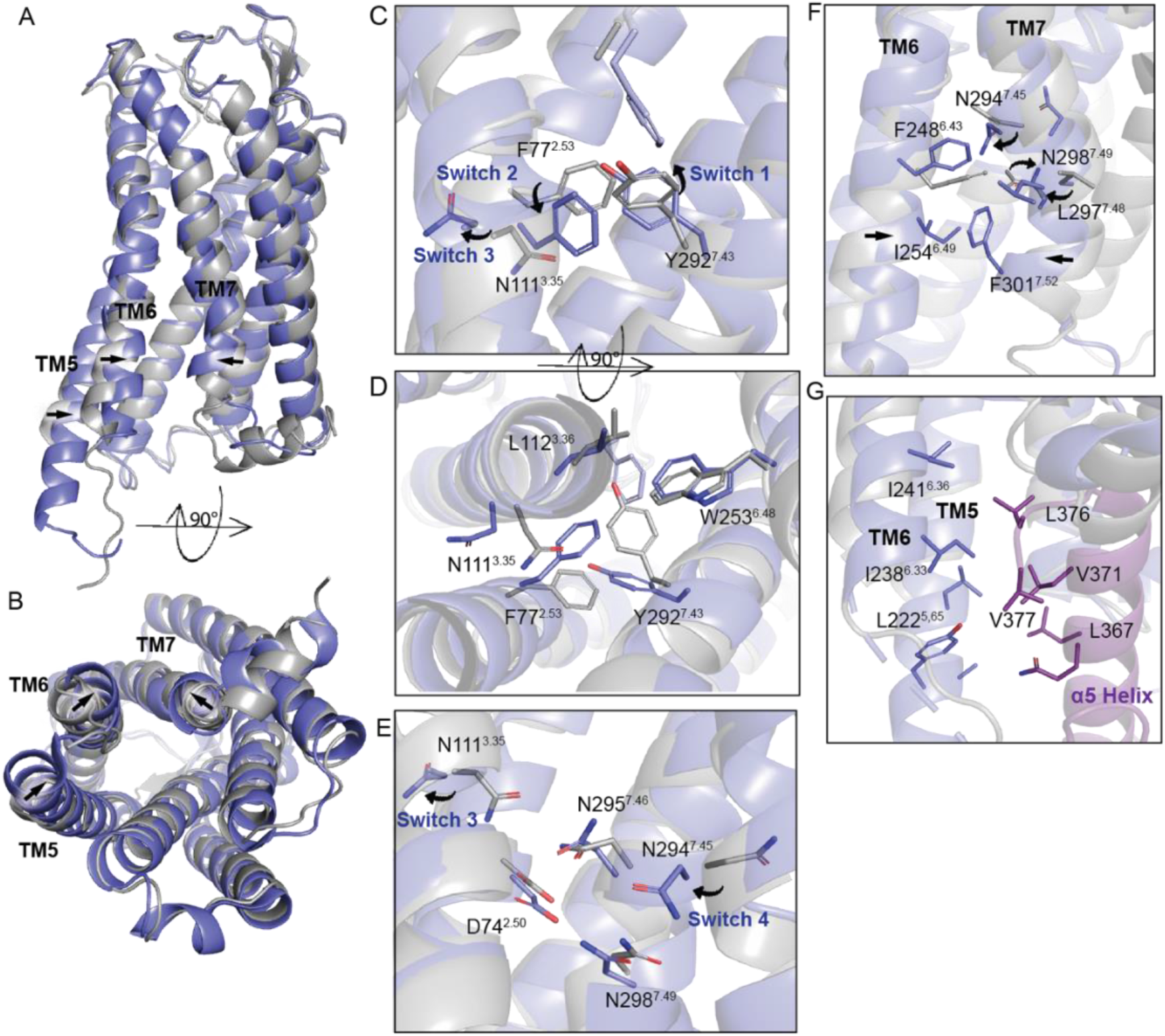
Structure comparisons between TRV055-bound AT1R and AT110-bound TRV023-bound AT1R (6OS1). (A, B) Overall structural superposition of the TRV055-bound G protein-coupled AT1R structure (blue) and the previously reported TRV023-bound nanobody-stabilized structure (gray) viewed from the side (A) intracellular bottom (B). Arrows indicate shifts of the TMs. (C-E) Comparison of the G protein-coupled TRV055-bound structure (blue) with the previously reported TRV023-bound nanobody-stabilized AT1R structure (gray) at the orthosteric pocket region from side view (C) and bottom view (D), and at at lower TM region (E). Arrows indicate conformation rearrangements of the residues. Microswitches are labeled in dark blue. (F) TM6-TM7 interface comparisons between the two structures. Arrows indicate rearrangements of the residues at the interface and shifts of the two helices. (G) Binding interface between TRV055-bound AT1R (blue) and Gq α5 helix (purple).

Unlike TRV055 and AngII, TRV023 lacks the C-terminal Phe8 residue and therefore does not extend as deeply into the receptor core toward TM7. Comparing to TRV026-bound AT1R structure, in the TRV055-bound structure, the ligand C-terminus adopts a TM7-directed orientation that stabilizes Y292^7.43^ (switch 1) and promotes coordinated rearrangements involving F77^2.53^ (switch 2) and N111^3.35^ (switch 3) within the receptor core. The rotation of N111^3.35^ away from the receptor core disrupts the inactive/β-arrestin bound state-stabilizing interaction between N111^3.35^ and N295^7.46^, subsequently disrupting the restrictions between TM3 and TM7 (Fig. 4C, 4D).

Additional differences are observed in the lower TM7 region. As mentioned previously, in the TRV055-bound structure, stabilization of Y292^7.43^ alters the geometry of TM7, positioning N294^7.45^ (switch 4) inward toward the receptor core, where it forms a hydrogen-bonding interaction with N298^7.49^ in the NPxxY motif (Fig. 4E). This rearrangement also reorients hydrophobic residues at the TM6–TM7 interface to face one another (Fig. 4F), bringing the two helices into closer proximity relative to the nanobody-stabilized TRV023 structure, promoting hydrophobic interactions between Gq α5 helix and TM5 and TM6 (Fig. 4G). These observations indicate a more compact lower TM region associated with the G protein-coupled state (Fig 4A, 4B). Interestingly, although the olmesartan-bound structure adopts a more closed TM6–TM7 conformation than the Gq-coupled structure (Fig S6 A-C), a similar inward-facing arrangement of these hydrophobic residues is observed at the TM6–TM7 interface (Fig. S6G). This suggests that the local hydrophobic packing is maintained despite differences in the relative positions of TM6 and TM7.

### Functional effects of AT1R microswitch mutations on AT1R signaling

To evaluate the functional role of the residue rearrangements observed in the G protein-coupled structures, we introduced mutations at residues involved in the proposed microswitch network, including F77^2.53^, N111^3.35^, L112^3.36^, Y292^7.43^, N294^7.45^, and N298^7.49^, and measured ligand binding, Gq dissociation and β-arrestin recruitment using BRET assays (Fig. 5A-5H). Radioligand competition binding showed that most mutations retained or increased affinity for AngII and TRV055, indicating that their effects on signaling were not simply attributable to impaired ligand binding (Fig. 5A, 5B, and S8). Most mutants also retained substantial Gq activation in response to either AngII or TRV055, indicating that these residues are not strictly required for receptor activation. In contrast, several mutations produced markedly stronger effects on β-arrestin recruitment and altered signaling bias profiles (Fig. 5C-H, and S8).

**Figure 5.**
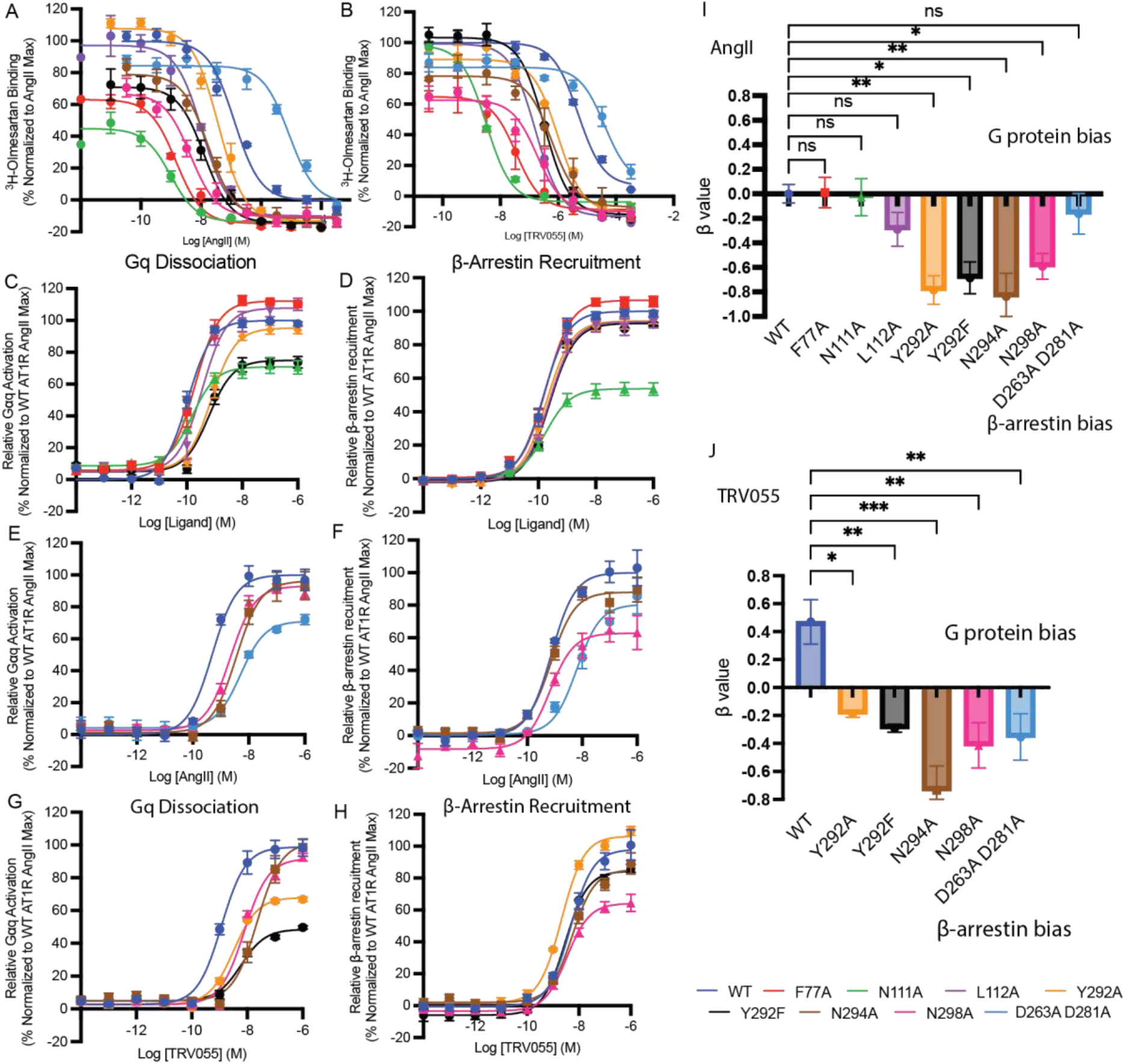
Binding and signaling bias of AngII and TRV055 across receptor mutants. (A,B) Competition binding between the radiolabeled antagonist [^3^H]-olmesartan and unlabeled AngII (A) and TRV055 (B) in cell membranes containing AT1R mutants. (C-F) BRET-based Gq dissociation (C, E) and β-arrestin recruitment (D, F) assays measuring AngII-induced signaling by wild-type (WT) AT1R and the indicated receptor mutants. (G, H) BRET-based Gq dissociation (G) and β-arrestin recruitment (H) assays measuring TRV055-induced signaling by wild-type (WT) AT1R and the indicated receptor mutants. Expression data is shown in Fig. S8. (I) AngII and (J) TRV055 bias factors for each mutant calculated from Gq dissociation and β-arrestin recruitment assays. WT AT1R signaling induced by AngII was used as a reference for each mutant and ligand. Error bars represent the mean SE of at least 3 independent experiments. β values were presented as mean ± S.E.M. β > 0, Gq biased; β < 0, β-arrestin-biased. Statistics analyses were performed using one-way analysis of variance (ANOVA). P < 0.05 (*); P<0.01 (**).

Notably, mutations in the orthosteric pocket, including Y292A, and residues within the lower TM7 region, including N294A, and N298A, shifted AngII signaling toward a more β-arrestin-biased profile relative to wild-type AT1R, with bias factors of -0.784, -0.836, and -0.591, respectively. These mutations did not reduce AngII binding affinity, further supporting that the observed changes in signaling reflect altered receptor coupling rather than impaired ligand recognition. Consistent with these results, Y292A, N294A, and N298A were among the mutations previously identified in a comprehensive alanine-scanning study of AT1R as reducing Gq signaling and shifting signaling toward β-arrestin bias^21^. Interestingly, the same mutations also altered the signaling properties of TRV055, converting this normally G protein-biased ligand (bias factor 0.47) into a more β-arrestin-biased agonist with bias factors of -0.187 (Y292A), -0.735 (N294A), and -0.414 (N298A). Binding affinity for TRV055 was also largely preserved in these mutants, indicating that the loss of G protein bias was not driven by reduced ligand binding. Unexpectedly, mutation of Y292^7.43^ to phenylalanine also altered signaling bias (bias factors -0.685 for AngII and -0.291 for TRV055), despite preservation of the aromatic side chain and the absence of direct hydrogen-bonding interactions involving the Y292^7.43^ hydroxyl group in the structure. One possible explanation is that the polar hydroxyl group of Y292^7.43^ increases the size, polarity and conformational constraints of the side chain, thereby stabilizing a specific rotameric state required for proper TM7 packing.

Mutations within the receptor core, including F77A and L112A, produced more modest effects (Fig. 5G, 5H). An interesting observation was that the D263A/D281A double mutation substantially reduced binding affinity for both AngII and TRV055 but had different effects on their signaling bias. The mutation had little effect on AngII signaling but markedly reduced the G protein bias of TRV055 (Fig. 5G, 5H), shifting it toward β-arrestin-biased signaling. This differential effect may reflect distinct dependence of the two ligands on the electrostatic environment of the extracellular pocket. AngII may compensate for the loss of these acidic residues through its broader interaction network, including contacts mediated by Arg2 and the charged N-terminus Asp. By contrast, TRV055 lacks the Arg2 interaction and may depend more strongly on D263^6.58^ and D281^7.32^ to define the geometry of its N-terminal binding environment. Removing both residues may therefore increase local flexibility or alter the packing between TM6 and TM7, selectively destabilizing receptor conformations that support G protein activation while preserving β-arrestin signaling.

Together, these results support a model in which coordinated rearrangements surrounding Y292^7.43^ and the lower TM7 region play an important role in regulating signaling bias at AT1R independently of their effects on ligand binding affinity.

## Discussion

In this study, we determined cryo-EM structures of wild-type AT1R coupled to heterotrimeric miniGs/q/iN in complex with the endogenous agonist AngII and the G protein-biased agonist TRV055. Together with peptide analog and receptor mutagenesis studies, these structures provide structural insight into how subtle changes in ligand conformation are translated into coordinated rearrangements within the receptor core that regulate signaling bias. In particular, our results identify an extended microswitch network connecting the orthosteric pocket to the lower TM7 region and the intracellular G protein-binding interface, providing a structural framework for understanding G protein-biased activation of AT1R.

Biased signaling at AT1R has been extensively investigated as a strategy to improve cardiovascular therapeutics^4–6,11,22^. To date, structural studies have predominantly focused on balanced and β-arrestin-biased ligands^16,17^. However, the structural basis by which G protein-biased agonists preferentially stabilize G protein signaling has remained comparatively unexplored. Our structures address this gap by providing the first structural characterization of a G protein-biased AT1R agonist bound to a G protein. Despite differing from AngII by only subtle N-terminal sequence changes, TRV055 adopts a nearly identical binding mode while promoting coordinated receptor rearrangements associated with G protein activation, suggesting that G protein bias can arise from fine-tuning receptor conformation rather than large changes in ligand recognition.

The peptide analog analysis further highlights the importance of the N-terminal glycine in TRV055 for preferential G protein activation. Substitution of Gly with bulkier residues consistently reduced G protein bias, suggesting that the additional space created by glycine near D263^6.58^ and D281^7.32^ contributes to stabilization of receptor conformations favorable for G protein coupling. Notably, this observation is consistent with another G protein-biased AT1R peptide, TRV056 (DRGVYIHPF)^23^, which similarly contains a glycine inserted at the position corresponding to Arg2 in AngII. Although the precise molecular mechanism remains to be established, the shared presence of glycine in two independently G protein-biased ligands supports the notion that increased conformational space at the ligand N-terminus may be a common structural feature promoting preferential G protein signaling.

The structural basis of AT1R biased signaling has previously been investigated using complementary spectroscopic and computational approaches^18,24^. DEER spectroscopy demonstrated that ligands with distinct signaling profiles stabilize different intracellular conformational ensembles, with G protein-biased ligands preferentially populating more open intracellular conformations and β-arrestin-biased ligands favoring comparatively occluded states^24^. Molecular dynamics simulations subsequently proposed that these distinct ensembles arise from conformational dynamics of the ligand C-terminal F8, which samples vertical, horizontal, and inserted conformations^18^. In particular, the simulations suggested that the inserted conformation enables direct interaction between F8 and Y292^7.43^, helping AngII and G protein-biased ligands stabilize the canonical active TM7 conformation. Our structures provide direct structural support for this hypothesis by showing that both AngII and TRV055 adopt a well-defined TM7-directed F8 conformation in the G protein-coupled state without detectable conformational heterogeneity. More importantly, our structures reveal how stabilization of this ligand conformation propagates through coordinated rearrangements of Y292^7.43^, F77^2.53^, N111^3.35^, L112^3.36^, N294^7.45^, and N298^7.49^, defining an allosteric pathway linking ligand recognition to remodeling of the intracellular G protein-binding interface.

Compared with the arrestin-mimetic nanobody-bound TRV023 structure, our G protein-coupled complexes exhibit a more compact TM6/TM7 interface, resulting from coordinated rearrangements within the TM7 region and the formation of a more tightly packed intracellular receptor core. Although the intracellular conformations observed in our structures differ from those inferred by DEER spectroscopy^24^, these observations likely reflect different stages of receptor activation rather than conflicting mechanisms. The DEER and molecular dynamics studies examined ligand-bound receptors in the absence of intracellular effectors and therefore described the intrinsic conformation available for effector coupling. By contrast, our structures capture effector-bound receptor complexes. A similar phenomenon has recently been observed for the serotonin 5-HT2A receptor, in which the Gq -bound structure adopts a narrower intracellular cavity than the β-arrestin-bound structure^25^. Together, these findings support a model in which ligand binding first reshapes the conformational landscape of AT1R, followed by additional structural remodeling upon engagement of G proteins or β-arrestins that stabilizes distinct effector-specific active conformations.

Our functional results are consistent with the comprehensive alanine-scanning study by Cao *et al.*, which combined systematic mutagenesis with molecular dynamics simulations to map the allosteric communication network governing Gαq and β-arrestin coupling at AT1R^21^. That study showed that G protein signaling depends on a broad network of residues distributed throughout the receptor, whereas β-arrestin coupling relies on fewer structural determinants, and identified extensive allosteric communication extending from the orthosteric pocket to the intracellular coupling interface. Among these, mutations at Y292^7.43^, N294^7.45^, and N298^7.49^ were also reported to shift receptor signaling toward β-arrestin bias. We independently observed similar functional effects, with mutation of these same residues shifting AngII toward β-arrestin bias and, notably, converting the G protein-biased agonist TRV055 into a β-arrestin-biased ligand. The agreement between these independent studies further supports an important role for the TM7 region in regulating signaling bias at AT1R.

Collectively, our work bridges previous spectroscopic observations, computational predictions, and functional studies by providing complete structural view of G protein coupling to wild-type AT1R, from ligand recognition in the orthosteric pocket through receptor microswitch rearrangements to the intracellular G protein interface. These findings establish a mechanistic framework linking ligand recognition, receptor activation, and transducer coupling, and provide a structural basis for the rational design of biased AT1R therapeutics.

## Limitations of the study

This study has several limitations. First, our cryo-EM structures represent static snapshots of the G protein-coupled state and therefore do not capture the dynamic conformational transitions underlying biased signaling. Although previous molecular dynamics simulations have provided important insight into ligand-dependent conformational changes, these studies were performed in the absence of intracellular effectors, whereas our structures capture only the effector-bound end state. Future simulations incorporating the G protein will therefore be valuable for defining how ligand-induced conformational ensembles evolve into fully coupled signaling complexes. Second, our structural comparisons rely on a G protein-bound AngII/TRV055 complex and a previously reported arrestin-mimetic nanobody-stabilized TRV023 structure. Determination of β-arrestin-bound AT1R structures, particularly in comparison with the same ligand bound to different effectors, will be important for defining the complete structural trajectory underlying biased signaling.

## Data Availability

The cryo-EM models and maps were deposited under the following accession numbers: Structure of G protein-coupled angiotensin II type I receptor bound to TRV055 PDB: 37YZ, EMDB: EMD-78654; Structure of G protein-coupled angiotensin II type I receptor bound to Angiotensin II PDB: 37YQ, EMDB: EMD-78648. All other data are included in the manuscript and/or SI Appendix.

## Acknowledgments

All cryo-electron microscopy data were collected at the Harvard Center for Cryo-Electron Microscopy at Harvard Medical School, and we thank the staff for assistance and advice during data collection. The SBGrid Consortium provided computational Support for structural biology software. This work was funded by a Merck Fellowship from the Helen Hay Whitney Foundation (M.A.S.), R01CA260415 (A.C.K.), R21HD101596 (A.C.K.), and support from the Smith Family Foundation (A.C.K.).

## Author Contributions

J.L., M.A.S., and A.C.K. designed research; J.L. and M.A.S. performed research; J.L. and A.C.K. analyzed data; and J.L. wrote the paper with input from all authors.

## Materials and Methods

### Constructs

### AT1R-miniGαs/qiN expression and purification

Expi293F TetR cells were grown in Expi293 media and transfected with the Flag-AT1R-miniGαs/qiN fusion protein^20^ (750 μg / L of culture) with Fectopro (800 μL/ L of culture). The culture was supplemented with 3 mM valproic acid and 0.4% (w/v) glucose 18 hours after transfection and cultured for 24 more hours. Receptor expression was then induced with 0.4 μg / mL doxycycline and the culture was supplemented with 5 mM sodium butyrate. Cell pellets were harvested 30 hours post induction. Cell pellets were thawed and lysed by resuspension in hypotonic buffer (10 mL of 10 mM Tris pH 7.4, 2 mM EDTA, 10 mM MgCl_2_ / g of cell pellet), with benzonase nuclease (Sigma Aldrich) and Pierce protease inhibitor tablets (ThermoFisher). Cells were pelleted (50,000 x g, 15 min) and resuspended in cold solubilization buffer 20 mM HEPES pH 7.4, 500 mM NaCl, 0.5% LMNG (w/v), 0.05% CHS (w/v), 10 mM MgCl_2_, benzonase nuclease, Pierce protease inhibitor tablets) via Dounce homogenization. The cell mixture was stirred for 2 hr at 4 °C to solubilize the membranes and insoluble material was pelleted by centrifugation (50,000 x g, 30 min). The supernatant was supplemented with 2 mM CaCl_2_, filtered through a glass fiber filter, and passed over anti-FLAG M1 resin. The resin was washed with 20 column volumes of LMNG wash buffer (20 mM HEPES pH 7.4, 500 mM NaCl, 0.01% LMNG (w/v), 0.01% CHS (w/v), 2 mM CaCl2) and eluted with LMNG wash buffer supplemented with 5 mM EDTA and 0.2 mg/mL Flag peptide (DYKDDDDK) and with removed calcium. The receptor was concentrated using a a 50 kDa cutoff Amicon-Ultra centrifugal filter and loaded onto a Superdex S200 (10/300) Increase column in 20 mM HEPES pH 7.4, 100 mM NaCl, 0.01% LMNG (w/v), 0.001% CHS (w/v).

### Gβγ expression and purification

Gβγ proteins were expressed and purified as previously described with modifications^26^. Briefly, *Trichuplusia ni* (Hi5) insect cells (Expression Systems) were infected with virus encoding the wild-type human β1γ2 subunits with a decahistidine tag inserted at the amino terminus of the β1 subunit with HRV-3C protease cleavable site. Cells were harvested 48 h post infection, lysed in hypotonic buffer (10 mM Tris pH 7.4, 5 mM β-ME and protease inhibitors) and Gβγ protein was extracted in a buffer containing 20 mM HEPES pH 7.5, 100 mM NaCl, 1% sodium cholate (w/v) (Sigma-Aldrich), 0.05% DDM (w/v), 5 mM β-ME, protease inhibitors, and 5 mM imidazole. The cell mixture was stirred for45 mins at 4 °C and insoluble material was pelleted by centrifugation (50,000 x g, 20 min). Clarified lysate was loaded onto a 5 mL Ni-NTA affinity column (HiTrap FF, Cytiva) equilibrated with solubilization buffer. Following binding, detergent was gradually exchanged to 0.1% DDM (w/v) by sequential washes containing decreasing proportions of solubilization buffer (50%, 25%, 10%, and 5%), followed by washing with buffer containing 20 mM HEPES (pH 7.5), 100 mM NaCl, 0.1% DDM (w/v), 1 mM MgCl₂, 5 mM β-mercaptoethanol (β-ME), and 20 mM imidazole. Protein was eluted with the same buffer supplemented with 400 mM imidazole. After elution, HRV-3C protease was added and the protein was dialyzed against a buffer containing 20 mM HEPES pH 7.5, 100 mM NaCl, 1 mM MgCl_2_, 0.05% DDM (w/v), 100 μM Tris(2-carboxyethyl)phosphine hydrochloride (TCEP) (Sigma-Aldrich), 20 mM GDP (Sigma-Aldrich). Cleaved heterotrimeric G-protein was further purified by reloading over Ni-sepharose resin. The flow through was collected and was treated with λ phosphatase (New England Biolabs), calf intestinal phosphatase (CIP) (ThermoFisher), and Antarctic phosphatase (New England Biolabs) in the presence of 1 mM MnCl₂ for 45 min on ice. The protein was further purified by anion-exchange chromatography on a MonoQ column (Cytiva) equilibrated in 20 mM HEPES (pH 7.5), 50 mM NaCl, 0.02% DDM (w/v), and 100 μM TCEP. After washing, bound protein was eluted using a linear NaCl gradient (50–500 mM). Fractions corresponding to the fully geranylgeranylated Gβγ heterodimer were pooled, dialyzed against 20 mM HEPES (pH 7.5), 100 mM NaCl, 0.02% DDM (w/v), and 100 μM TCEP.

### ScFv16 expression and purification

ScFv16 was expressed and purified as described^27^ with modifications. Sf9 cells were grown in ESF 921 media at a density of 3 x 10^6^ cells/mL and infected with 5 mL virus / L of culture. After 48 hours, the supernatant was harvested (4000 x g, 15 min) and filtered through a 0.2um filter. 25 mM Tris pH 8.0 was added to the media to reach pH 7.5 and 1mM nickel chloride and 5mM calcium chloride were added to precipitate the chelators. The mixture was stirred for 1 hour at room temperature. Resulting precipitates were removed by centrifugation and the supernatant was passed over preequilibrated Ni-NTA resin. The column was washed with a high salt buffer (20 mM HEPES pH 7.4, 500 mM NaCl and 10 mM imidazole) followed by a low salt buffer (20 mM HEPES pH 7.5, 100 mM NaCl and 10 mM imidazole). The protein was eluted with the elution buffer (20 mM HEPES pH 7.5, 100 mM NaCl and 300 mM imidazole). The eluted protein was further purified over a size-exclusion chromatography using a Superdex 200 Increase 10/300 GL column (Cytiva). Monomeric fractions were pooled, concentrated and flash frozen in liquid nitrogen until use.

### Cryo-EM complex formation and sample preparation

The AT1R-miniGs/qiN fusion protein was mixed a 1.5-fold molar excess of the Gβγ and 2-fold molar excess of the ScFv16 and incubated on ice for 1.5 hour in the presence of 50 uM AngII (Sigma) or TRV055. Then 100 mU/ml Apyrase (Sigma) was added and then mixture was incubated on ice for another 1.5 hour. The sample was loaded onto anti-FLAG M1 resin, and washed with 6 column volumes of 20 mM HEPES pH 7.4, 100 mM NaCl, 0.01% LMNG (w/v), 0.001% CHS (w/v), 2 mM CaCl_2_ with of 50 μM of AngII or TRV055. The sample was then exchanged into glycol-diosgenin (GDN) detergent by washing with 20 column volumes of GDN wash buffer (20 mM HEPES pH 7.4, 100 mM NaCl, 0.01% GDN (w/v), 0.001% CHS (w/v), 2 mM CaCl_2_) supplemented with of 10 uM AngII or TRV055 and eluted with GDN wash buffer supplemented with with 5 mM EDTA and 0.2 mg/mL FLAG peptide. The complex was concentrated in a 100 kDa cutoff Amicon-Ultra centrifugal filter, and further purified by size exclusion chromatography (Superdex S200 Increase 10/300) in GDN wash buffer supplemented with 20 μM AngII or TRV055. Peak fractions were concentrated to 3-5 mg/mL. 3 μL of sample was applied to glow-discharged UltraAuFoil 300 mesh grids with 1.2 μm diameter /1.3 μm spacing (TedPella) at 20 °C with 100% humidity. Grids were blotted for 4-6 seconds with a blot force of 15 on a Vitrobot Mark IV Vitrobot (ThermoFisher) and then plunged into liquid-nitrogen-cooled liquid ethane.

### Cryo-EM data collection, and structure determination

Cryo-EM data were collected at the Harvard Cryo-Electron Microscopy Center for Structural Biology on a 300 kV Titan Krios G3i microscope in counted mode with a K3 detector GIF quantum energy Filter (20 eV) (Gatan) for TRV055-bound structure, and a 200 kV Talos Arctica with Volta phase plate and Gatan K3 direct electron detector for the AngII-bound structure. For the TRV055-bound structure, two movies were collected from the center of each hole in a nine-hole pattern per stage position with SerialEM data collection software v4.05^28^. For the AngII-bound structure, one movie was collected from the center of each hole in a nine-hole pattern per stage position with SerialEM data collection software v4.05. Collection parameters are listed in Table S1. Data processing was carried out in CryoSparc v4^29^ (Figs. S1 and S2). Movies underwent patch motion correction, dose-weighting, and patch contrast transfer function (CTF) estimation. Particles were picked with the blob-picker function and processed through 2D classification to identify a subset of particles to generate ab initio reconstructions. All particles were then classified through multiple rounds of heterogenous refinement and subjected to non-uniform refinement. The model of the complex was built in Coot^30^ and refined with Phenix^31^ real space refinement. In regions with density for the protein backbone, but weak side chain density, side chains were modeled with the most common rotamer position that did not clash with the model. Local resolution estimates were calculated in Phenix^31^. Figures were prepared in UCSF ChimeraX v1.6.1 ^32^and Pymol v3.1.3^32,33^. All structural biology software was supported by SBGrid^34^.

### Bioluminescence resonance energy transfer (BRET) assays

Dissociation of the Gq heterotrimer was measured using the G protein-based, tricistronic activity sensors (G-CASE)^35^. HEK293T cells were grown to 50% confluency in DMEM + 10% FBS + 1% penicillin-streptomycin and transfected with 50 ng of the Gq-CASE sensor and 750 ng of FLAG-AT1R plasmid with FugeneHD (3 μL/ μg DNA) in a 35 mm dish. 18 hours after transfection, cells were lifted with trypsin and 4E5 cells were plated per well in a clear-bottom, white-walled, poly-D-lysine coated 96 well plate in phenol red free, glucose free, and glutamine free DMEM + 2% FBS + 1x glutamax + 1% antibiotic-antimycotic. The next day, media was aspirated and replaced with 90 μL Hanks’ Balanced Salt Solution (HBSS) without calcium and magnesium supplemented with 20 mM HEPES pH 7.4 and 5 μμ coelenterazine-h. Cells were incubated for 5 min and baseline BRET ratios were read on a GloMax plate reader (Promega). Ligands were applied and BRET ratios were measured every 5 min for 15 min. Net BRET values were calculated by subtracting the baseline BRET ratio for each well followed by the ligand-free reading.

β-arrestin2 recruitment to AT1R was measured by BRET between nLuc β-arrestin2 and FLAG-AT1R-mVenus. HEK293T cells were grown to 50% confluency in DMEM + 10% FBS + 1% pen/strep and transfected with 5 ng of nLuc β-arrestin2 plasmid and 250 ng of FLAG-AT1R-mVenus plasmid with FugeneHD (3 μL/ μg DNA) in a 35 mm dish. 18 hours after transfection, cells were lifted with trypsin and 4 x 10^4^ cells were plated in poly-D-lysine coated 96 well clear bottom plates in phenol red free, glucose free, and glutamine free DMEM + 2% FBS + 1x glutamax + 1% antibiotic-antimycotic. The next day, media was aspirated and replaced with 90 μL Hanks’ Balanced Salt Solution (HBSS) without calcium and magnesium supplemented with 20 mM HEPES pH 7.4 and 5 μμ coelenterazine-h. After five minutes of incubation, baseline BRET ratios were read on a GloMax plate reader (Promega). Ligands were applied and BRET ratios were measured every 5 min for 15 minutes. Net BRET values were calculated by subtracting the baseline BRET ratio for each well followed by the ligand-free reading.

### Radioligand competition binding assays

Expi293F cells were transiently transfected with FLAG-AT1R containing plasmids with Fectopro. The culture was enhanced with 3 mM valproic acid and 0.4% glucose after 18 hours. Receptor expression was induced with 0.4 mg/mL doxycycline hyclate two days post-transfection for approximately 24 hours. Cells were pelleted, washed with cold HBS (20 mM Hepes pH 7.4, 150 mM NaCl), and resuspended in 2.5 mL of 20 mM Tris pH 7.4 per gram of cell pellet with a protease inhibitor tablet. Cells were lysed by Dounce homogenization (100x) and membranes were pelleted by centrifugation (50,000 xg for 20 min). Isolated membranes were resuspended in radioligand binding buffer (50 mM Tris pH 7.4, 12.5 mM MgCl_2_, 150 mM NaCl, 0.2% BSA) with a protease inhibitor tablet, flash frozen in liquid nitrogen, and stored at -80°C.

Cell membranes were thawed, resuspended in radioligand binding buffer, and incubated with varying amounts of cold ligands and 2 nM [^3^H]-olmesartan (American Radiolabeled Chemicals) in 200 mL reaction volumes for 90 min at room temperature. Reactions were harvested on a 96-well Brandel harvester with a GF/B filter soaked in water and washed three times with cold water. Data were fit with a one-site competition binding model in GraphPad Prism. [^3^H]-olmesartan affinity (K_d_) was determined for each variant through saturation binding assays by varying the amount of radioligand to calculate inhibitory constant (K_i_) values from competition binding experiments. Saturation binding data was fit with the one-site saturation binding model with non-specific binding of [^3^H]-olmesartan, measured in the presence of 10 mM candesartan, taken into account.

### Quantifying cell surface expression

Surface expression levels of AT1R mutants were determined using flow cytometry from cells prepared for BRET-based signaling assays. For Gαq dissociation assays, surface levels of FLAG-AT1R were monitored via M1 αFLAG AF647. For β-arrestin2 recruitment surface levels of FLAG-AT1R-mVenus were monitored via M1 αFLAG AF647 and total AT1R levels were detected via the C-terminal mVenus. Cells in a 6 well plate were washed with PBS (137mM NaCl, 2.7mM KCl, 10mM NaPO_4_, 1.8mM KPO_4_, pH 7.4), resuspended in 400 μL PBS and incubated with shaking at 60 rpm in 4 °C for 10 minutes to detach. 200 μL of dissociated cells from each well was plated in a 96 v-well bottom plate and washed with flow cytometry buffer (20 mM HEPES pH 7.5, 150 mM NaCl, 0.1% BSA, 2 mM CaCl_2_). Cells were stained in flow cytometry buffer supplemented with 100 nM M1α-FLAG AF647 for 20 min with shaking at 30 rpm at 4 °C. After incubation, wells were washed once with flow cytometry buffer and resuspended in 50 μL flow cytometry buffer for analysis on a CytoFLEX flow cytometer (Beckman Coulter). Relative expression values of mutants were calculated by normalizing to the wild type AT1R transfected condition as 100% expression and empty vector transfected condition as 0% expression.

### Bias factor calculation

Data from BRET assay were used to calculate the β values of different mutations using the following equation, where P1 represents Gq signaling and P2 represents β-arrestin recruitment. AngII signaling from WT AT1R was used as a reference for each ligand. Data were analyzed by GraphPad Prism 9.0 and presented as Mean ± S.E.M. from three independent experiments. Statistical significance was evaluated by one-way ANOVA test.

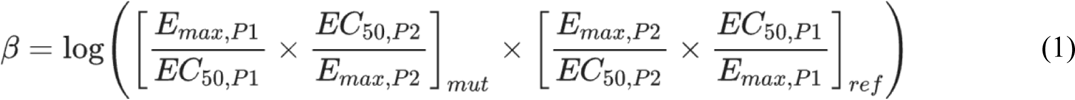

