## Supplementary information for "Structural Basis for Angiotensin II Type I Receptor G Protein-biased Signaling"

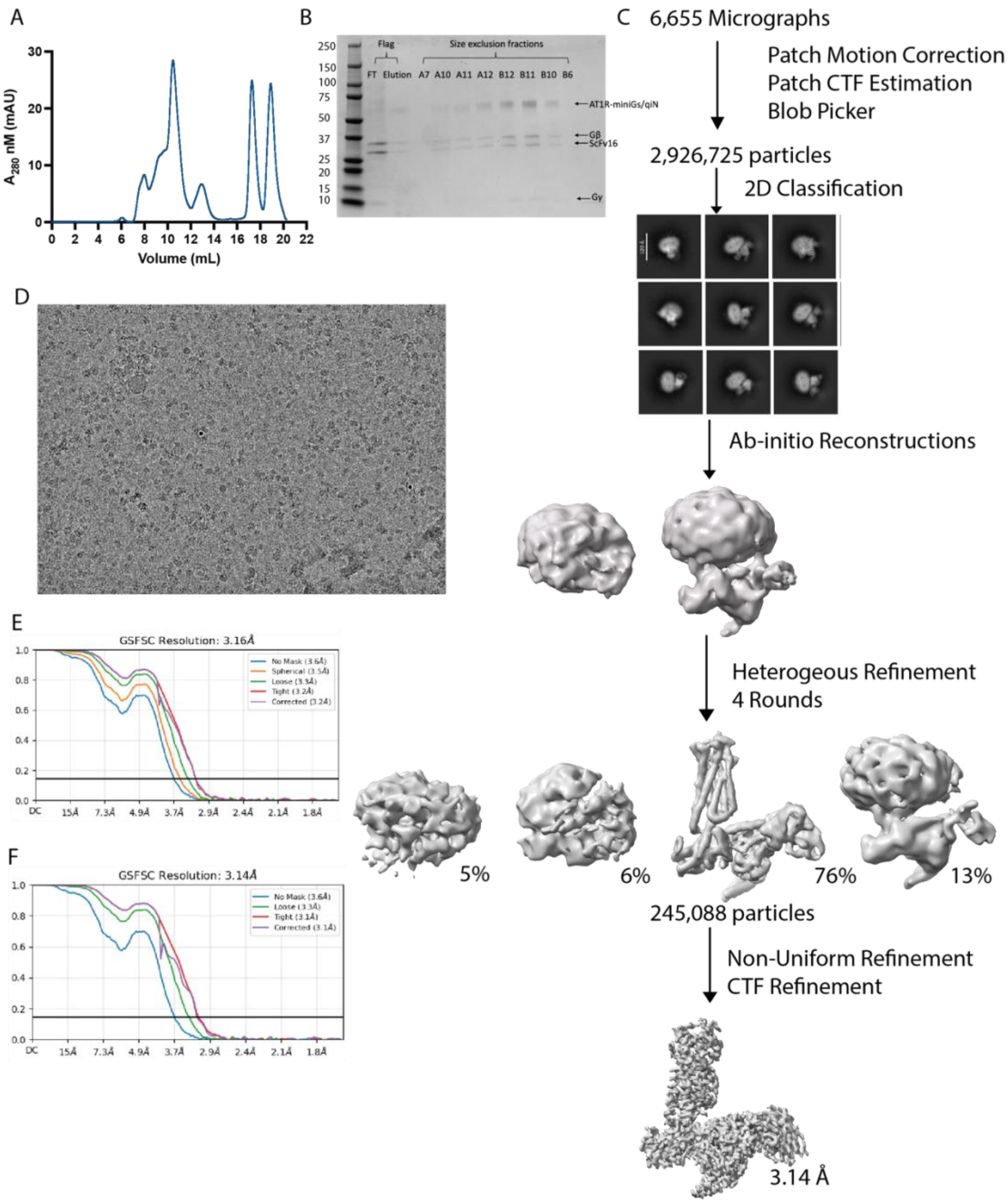

Figure S1. TRV055-bound AT1R-Gq complex data processing. (A) Size exclusion trace, (B) SDS-PAGE gel, (C) cryo-EM data processing scheme, and (D) representative micrograph. Fourier shell correlation (FSC) used to determine E) global map and F) locally refined map resolution.

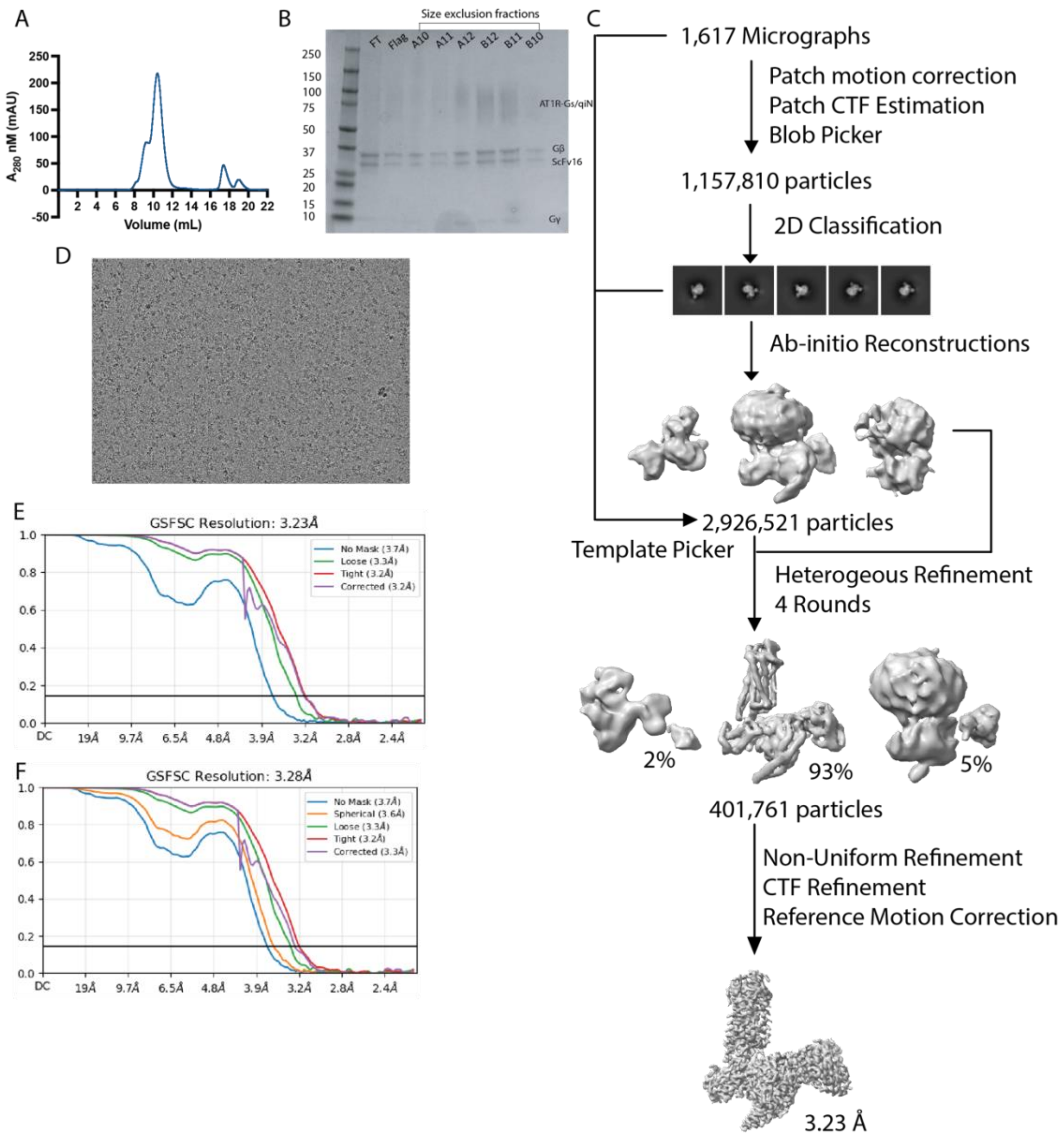

Figure S2. AngII-bound AT1R-Gq complex data processing. (A) Size exclusion trace, (B) SDS-PAGE gel, (C) cryo-EM data processing scheme, and (D) representative micrograph. Fourier shell correlation (FSC) used to determine E) global map and F) locally refined map resolution.

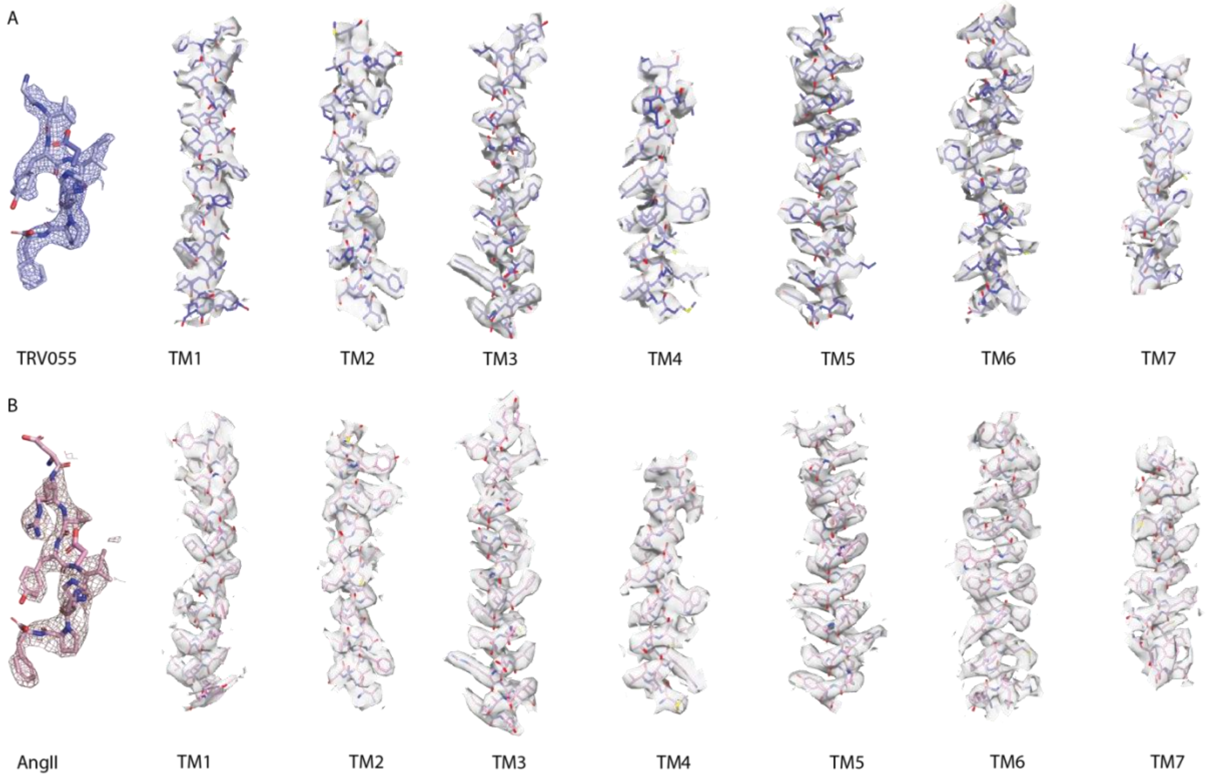

Figure S3. Cryo-EM density maps of the ligands and transmembrane helices. (A) Maps for TRV055 and TM1-7 for TRV055-bound AT1R structure. (B) Maps for AngII and TM1-7 for AngII-bound AT1R structure.

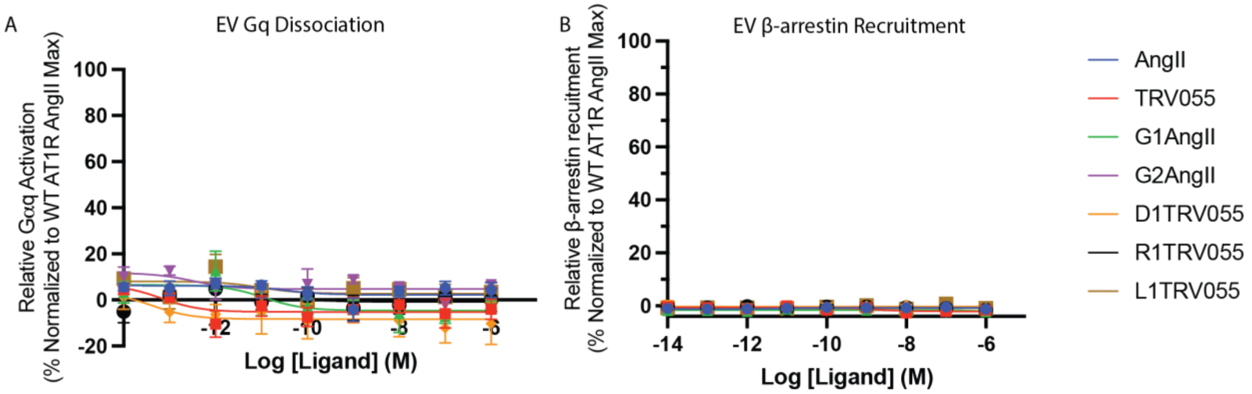

C

| | | | Gq Dissciation | | | | β-arrestin Recruitment | | | | |
|---|---|---|---|---|---|---|---|---|---|---|---|
| | LogKi (SEM) | | LogEC50 (SEM) | | Emax (SEM) | | LogEC50 (SEM) | | Emax (SEM) | | Bias Factor |
| AngII | -7.76 | 0.09 | -9.43 | 0.04 | 100.00 | 1.53 | -9.44 | 0.03 | 100.00 | 1.30 | 0.00 |
| TRV055 | -6.17 | 0.06 | -8.76 | 0.07 | 104.03 | 3.15 | -8.39 | 0.05 | 96.42 | 1.87 | 0.467 ± 0.104 |
| G1AngII | -8.64 | 0.08 | -9.61 | 0.12 | 104.80 | 4.92 | -9.53 | 0.09 | 98.63 | 3.39 | 0.118 ± 0.165 |
| G2AngII | -5.59 | 0.10 | -8.20 | 0.09 | 108.28 | 4.30 | -7.91 | 0.06 | 82.66 | 2.31 | 0.423 ± 0.123 |
| D1TRV055 | -4.96 | 0.15 | -6.93 | 0.15 | 96.35 | 8.64 | -6.86 | 0.08 | 88.59 | 4.23 | 0.129 ± 0.179 |
| R1TRV055 | -6.49 | 0.07 | -9.46 | 0.11 | 104.26 | 4.49 | -9.27 | 0.07 | 93.95 | 2.60 | 0.248 ± 0.146 |
| L1TRV055 | -5.05 | 0.13 | -7.44 | 0.12 | 84.87 | 5.01 | -7.36 | 0.04 | 83.01 | 1.79 | 0.107 ± 0.143 |

Figure S4. BRET-based Gq dissociation (A) and β-arrestin recruitment (B) assays performed in cells transfected with empty vector. AngII, TRV055, and the indicated peptide analogs did not elicit detectable signals over the concentration range tested. (C) Effects of AngII analogs on binding, Gq dissociation and β-Arrestin recruitment. Values are extracted from curve fits displayed in Fig. 2D, 2E. Data from three or more independent experiments are normalized relative to the %WT response to AngII.  Bias Factors were calculated use equation (1).

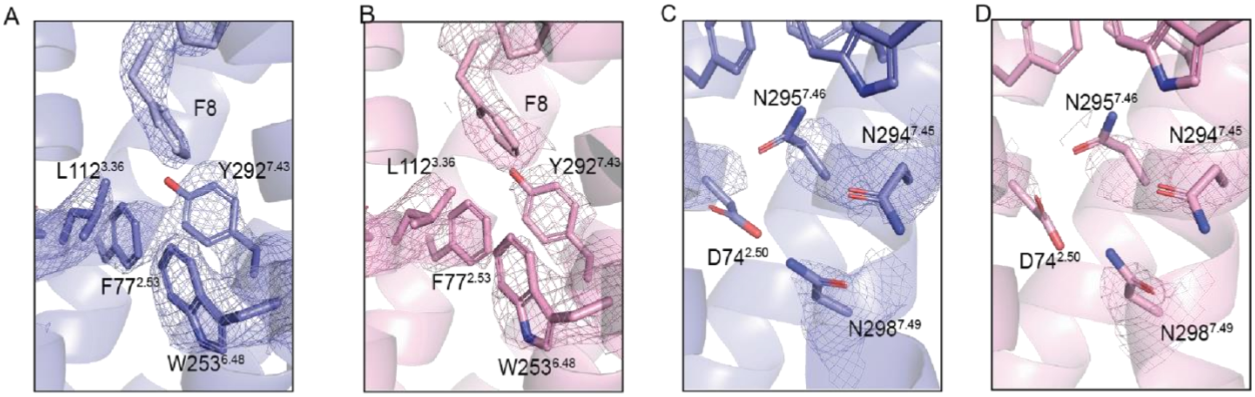

Figure S5. (A, B) Local cryo-EM map for the ligand C-terminal Phe8 and residues in the orthosteric pocket in the TRV055-bound (A) and AngII-bound (B) AT1R–Gq complexes. (C, D) Local cryo-EM map for the lower TM7 region in the TRV055-bound (C) and AngII-bound (D) AT1R–Gq complexes.

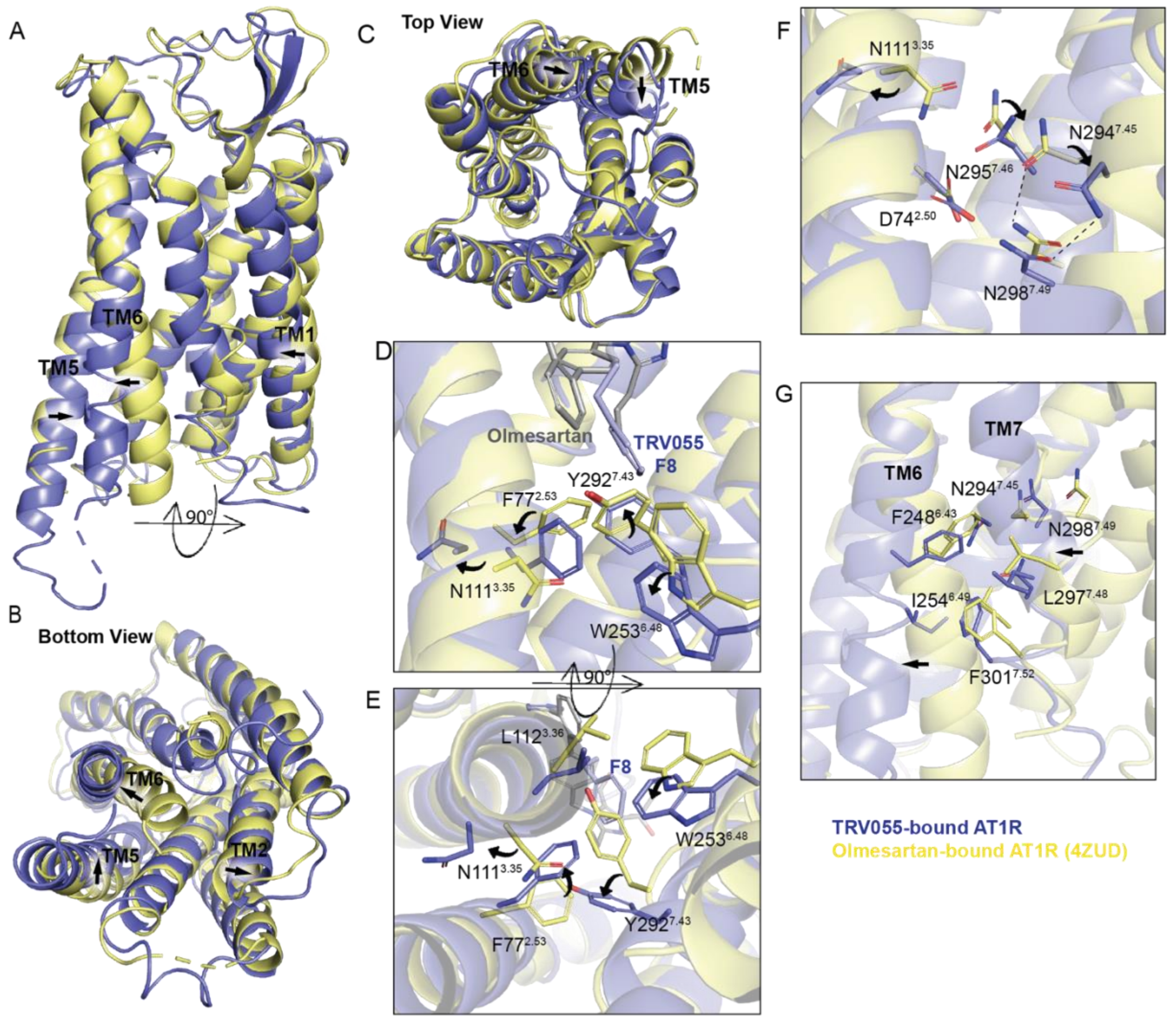

Figure S6. Structure comparisons between TRV055-bound AT1R and olmesartan-bound AT1R (4ZUD). (A-C) Overall structural superposition of the TRV055-bound G protein-coupled AT1R structure (blue) and olmesartan-bound AT1R structure (yellow) viewed from the side (A) intracellular bottom (B), and extracelluar top (C). Arrows indicate shifts of the TMs. (D-F) Comparison of the G protein-coupled TRV055-bound structure (blue) with olmesartan-bound AT1R structure (yellow) at the orthosteric pocket region from side view (D) and bottom view (E), and at at lower TM region (F). Arrows indicate conformation rearrangements of the residues. Microswitches are labeled in dark blue. (G) TM6-TM7 interface comparisons between the two structures. Arrows indicate rearrangements of the residues at the interface and shifts of the two helices.

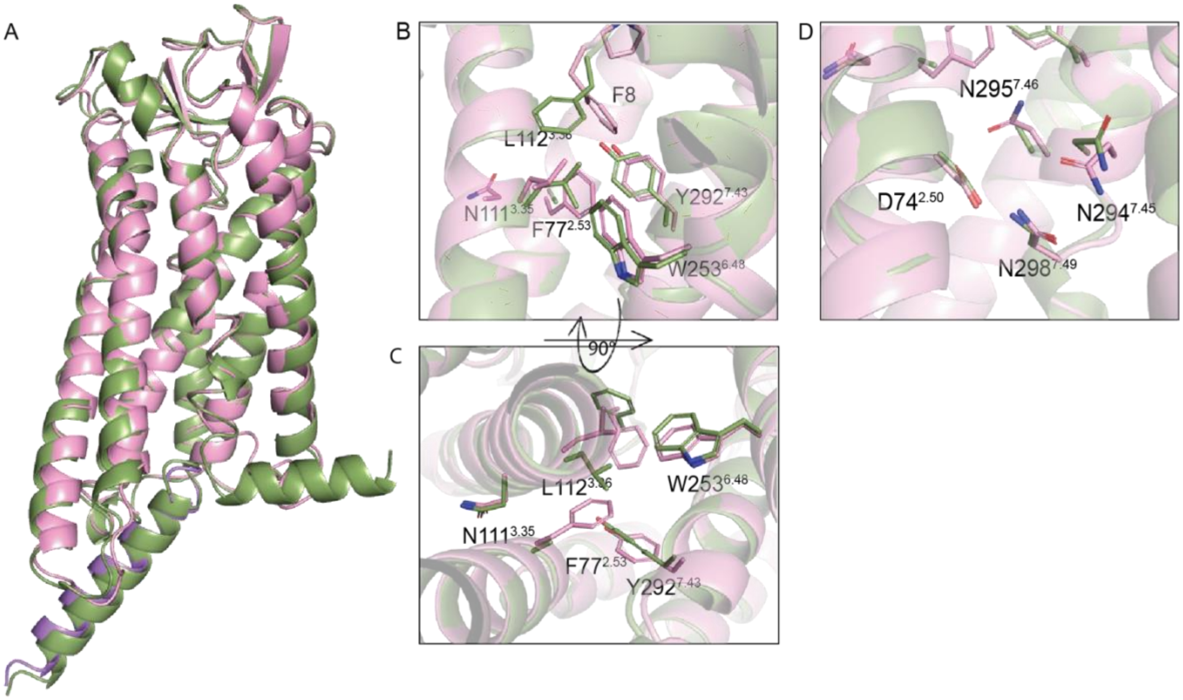

Figure S7. Structure comparisons between WT AngII-bound AT1R (pink) and Sar$^{1}$AngII-bound AT1R with F77A and N295A mutations (7F6G) (green). (A) Overall structural superposition of the two structures. (B-D) Comparison of the WT AT1R (pink) with the previously reported AT1R mutant (green) at the orthosteric pocket region from side view (B) and bottom view (C), and at at lower TM region (D).

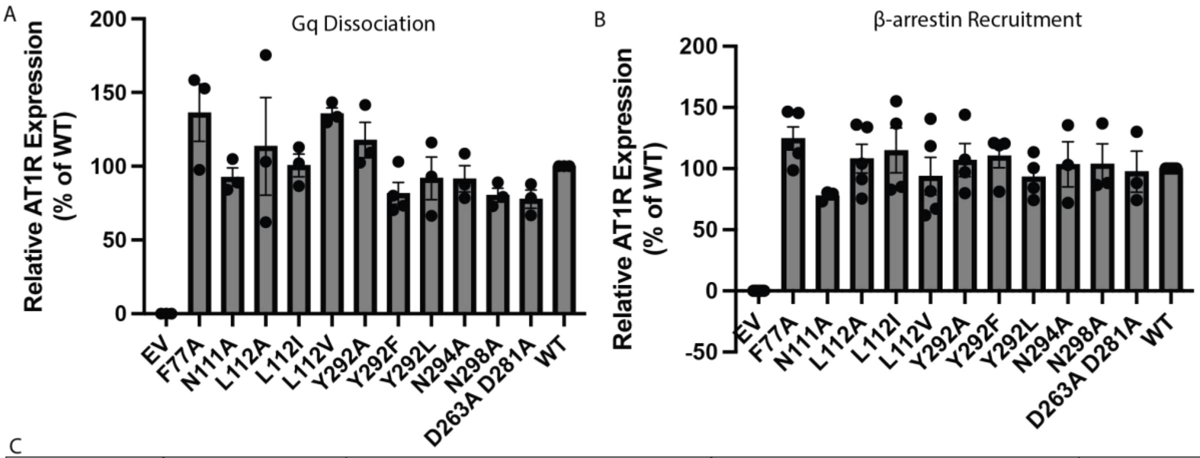

C

| | Competition Binding | | Gq DisscIation | | | | β-arrestin Recruitment | | | | |
|---|---|---|---|---|---|---|---|---|---|---|---|
| | LogKi (SEM) | | LogEC50 (SEM) | | Span (SEM) | | LogEC50 (SEM) | | Span (SEM) | | Bias Factor |
| WT | -7.76 | 0.09 | -9.97 | 0.06 | 100.00 | 2.39 | -9.81 | 0.05 | 100.00 | 2.01 | 0.00 |
| F77A | -9.15 | 0.09 | -9.71 | 0.07 | 106.38 | 2.85 | -9.74 | 0.05 | 105.90 | 2.12 | 0.0102 ± NA |
| N111A | -9.10 | 0.11 | -9.82 | 0.09 | 62.24 | 2.19 | -9.75 | 0.09 | 53.20 | 1.87 | -0.0272 ± 0.152 |
| L112A | -8.13 | 0.11 | -9.45 | 0.09 | 102.82 | 3.39 | -9.62 | 0.07 | 95.26 | 2.46 | -0.29 ± 0.138 |
| Y292A | -7.88 | 0.08 | -9.11 | 0.06 | 89.37 | 2.28 | -9.71 | 0.06 | 95.78 | 2.07 | -0.784 ± 0.116 |
| Y292F | -8.38 | 0.12 | -9.20 | 0.09 | 70.08 | 2.63 | -9.60 | 0.04 | 94.93 | 1.36 | -0.685 ± 0.13 |

D

| | Competition Binding | | Gq DisscIation | | | | β-arrestin Recruitment | | | | |
|---|---|---|---|---|---|---|---|---|---|---|---|
| | LogKi (SEM) | | LogEC50 (SEM) | | Span (SEM) | | LogEC50 (SEM) | | Span (SEM) | | Bias Factor |
| WT | -7.76 | 0.09 | -9.31 | 0.09 | 100.00 | 3.57 | -9.13 | 0.08 | 100.00 | 3.28 | 0.00 |
| N294A | -7.77 | 0.10 | -8.44 | 0.11 | 94.98 | 4.31 | -9.14 | 0.09 | 86.51 | 3.04 | -0.838 ± 0.19 |
| N298A | -8.36 | 0.13 | -8.67 | 0.07 | 90.24 | 2.66 | -9.18 | 0.14 | 71.02 | 4.07 | -0.591 ± 0.205 |
| D263A D281A | -5.24 | 0.12 | -8.30 | 0.07 | 66.84 | 1.89 | -8.20 | 0.08 | 79.70 | 2.95 | -0.163 ± 0.166 |

E

| | Competition Binding | | Gq DisscIation | | | | β-arrestin Recruitment | | | | |
|---|---|---|---|---|---|---|---|---|---|---|---|
| | LogKi (SEM) | | LogEC50 (SEM) | | Span (SEM) | | LogEC50 (SEM) | | Span (SEM) | | Bias Factor |
| WT | -6.17 | 0.06 | -8.94 | 0.05 | 106.40 | 2.34 | -8.33 | 0.08 | 98.24 | 3.38 | 0.47 ± 0.159 |
| F77A | -7.78 | 0.21 | N. D. | N. D. | N. D. | N. D. | N. D. | N. D. | N. D. | N. D. | N. D. |
| N111A | -8.58 | 0.05 | N. D. | N. D. | N. D. | N. D. | N. D. | N. D. | N. D. | N. D. | N. D. |
| L112A | -6.96 | 0.25 | N. D. | N. D. | N. D. | N. D. | N. D. | N. D. | N. D. | N. D. | N. D. |
| N294A | -6.22 | 0.19 | -7.64 | 0.09 | 96.70 | 4.14 | -8.26 | 0.08 | 83.23 | 2.80 | -0.735 ± 0.175 |
| N298A | -6.73 | 0.16 | -8.11 | 0.07 | 89.19 | 2.84 | -8.46 | 0.07 | 67.55 | 2.01 | -0.414 ± 0.162 |
| Y292A | -6.47 | 0.11 | -8.50 | 0.05 | 65.27 | 1.43 | -8.67 | 0.05 | 104.49 | 2.02 | -0.187 ± 0.0255 |
| Y292F | -6.96 | 0.06 | -8.22 | 0.08 | 43.72 | 1.58 | -8.54 | 0.06 | 90.40 | 2.00 | -0.291 ± 0.029 |
| D263A D281A | -4.55 | 0.14 | -8.08 | 0.07 | 74.56 | 2.37 | -8.22 | 0.08 | 79.55 | 2.86 | -0.354 ± 0.166 |

Figure S8. Surface expression levels, detected by AF647 conjugated M1-αFLAG in the Gq dissociation (A) and β-arrestin recruitment assays (B), were normalized across variants. Error bars represent the mean SE of at least 3 independent experiments. (C, D) Effects of AT1R variants on AngII competition binding, AngII-induced Gq Dissociation and β-arrestin recruitment. (E) Effects of AT1R variants on TRV055 competition binding, TRV055-induced Gq Dissociation and β-arrestin recruitment. Values are extracted from curve fits displayed in Fig. 5. Data from three or more independent experiments are normalized relative to the %WT response to AngII. Bias Factors were calculated use equation (1).

| | | |
|---|---|---|
| **Data collection and processing** | Gq-coupled AT1R bound TRV055 | Gq-coupled AT1R bound AngII |
| Voltage (kV) | 300 | 200 |
| Electron exposure (e–/Å$^2$) | 63 | 50 |
| Defocus range (μm) | -0.8 to -1.8 | -0.8 to -2.0 |
| Pixel size (Å) | 0.83 | 1.1 |
| Symmetry imposed | C1 | C1 |
| Initial particle images (no.) | 2,926,725 | 1,157,810 |
| Final particle images (no.) | 245,088 | 401,761 |
| Map resolution (Å)<br>FSC threshold | 3.14<br>0.143 | 3.23<br>0.143 |
| **Refinement** | | |
| Model resolution (Å) | 3.14 | 3.23 |
| Model composition<br>Non-hydrogen atoms<br>Protein residues<br>Ligands | <br>9036<br>1142<br>1 | <br>9075<br>1145<br>1 |
| R.m.s. deviations<br>Bond lengths (Å)<br>Bond angles (°) | <br>0.30<br>0.43 | <br>0.22<br>0.35 |
| Validation<br>MolProbity score<br>Clashscore<br>Poor rotamers (%) | <br>1.78<br>11<br>0.9 | <br>1.92<br>7<br>0.9 |
| Ramachandran plot<br>Favored (%)<br>Allowed (%)<br>Disallowed (%) | <br>96.5<br>3.4<br>0.1 | <br>94.6<br>5.3<br>0.1 |

Table S1. CryoEM data collection, refinement and validation statistics.
